# *Pgbd5* impacts neuronal differentiation and double strand breaks signaling in the developing cortex

**DOI:** 10.1101/2023.05.09.539730

**Authors:** Alessandro Simi, Federico Ansaloni, Luca Spagnoletti, Damiano Mangoni, Pierre Lau, Diego Vozzi, Luca Pandolfini, Devid Damiani, Remo Sanges, Stefano Gustincich

## Abstract

Transposable Element Derived 5 (*Pgbd5*) is an evolutionary conserved gene encoding an endonuclease predominantly expressed in the nervous system and known to drive oncogenic DNA rearrangements in childhood solid tumors. However, its physiological role in brain development has remained poorly understood.

Here we show that Pgbd5 is required for proper neuronal differentiation and radial migration during mouse corticogenesis. *In vivo* knockdown of Pgbd5 impairs neurogenesis and cortical layering without affecting cell viability. Transcriptomics analysis reveal upregulation of cell cycle-related genes and downregulation of genes involved in mitochondrial oxidative metabolism, ribosomal function and neuronal differentiation, including markers of neocortical layer identity. Mechanistically, Pgbd5 depletion leads to a reduction of visible endogenous DNA double-strand breaks (DSBs) in neural progenitors, supporting a role in genome plasticity during cortical development. Ultra-deep whole genome sequencing at E14.5 shows no evidence of Pgbd5-dependent somatic rearrangements.

Together, our findings identify Pgbd5 as a domesticated transposase essential for neurogenesis.

## INTRODUCTION

Transposable elements (TEs) are mobile genetic sequences that constitute a substantial fraction of eukaryotic genomes. Their mobilization is a major driver of genetic variability throughout evolution. TEs are broadly classified into two groups: retrotransposons (class I) and DNA transposons (class II). While retrotransposons propagate through RNA intermediates and reverse transcription, DNA transposons move via a “cut and paste” mechanism ^1–3^, mediated by a transposase enzyme that excises and reinserts the element into new genomic sites. During evolution, host genomes have domesticated transposases for various biological functions, as exemplified by RAG1/2, transib-derived recombinases essential for V(D)J recombination and adaptive immunity ^4^.

The *PiggyBac Transposable Element Derived 5* (*Pgbd5*) gene represents an ancient domestication of a piggyBac DNA transposon, dating back approximately 500 million years ^5,6^. *Pgbd5* is almost exclusively expressed in the brain and aberrantly in childhood solid tumors ^6–8^. It encodes a catalytically active endonuclease capable of mediating stereotypical cut-and-paste DNA transposition ^7^. In childhood solid tumors, pathological *Pgbd5* overexpression leads to widespread genomic rearrangements ^8^. However, its physiological role in neurons remains unknown.

During mammalian corticogenesis, neural progenitors cells (NPC), including radial glia (RG) cells in the ventricular zone (VZ) and intermediate progenitors (IP) in the subventricular zone (SVZ), undergo successive rounds of proliferation. These progenitors generate glutamatergic projection neurons that migrate radially across the intermediate zone (IZ) to populate the forming cortical plate (CP) in an inside-first, outside-last sequence ^9,10^. Early-born neurons form the deep layers (DL; layer 6 and 5), which project to subcortical targets (striatum, thalamus, and spinal cord), whereas late-born neurons populate the upper layers (UL; layers 4 and 2/3), establishing cortico-cortical callosal connections ^11,12^. Importantly, multiple physiological DNA double strand breaks (DSBs) occur in NPCs during the mitotic phase and in post-mitotic neurons ^13–15^ .

Recurrent DSBs clusters have been shown to control the expression of neurodevelopmental genes and repaired by mechanisms of non-homologous end joining (NHEJ) ^16^.

Given the highly specific expression of *Pgbd5* the brain and its deep evolutionary conservation ^6^, we hypothesized that this gene plays a key role in mammalian cerebral cortex development.

## RESULTS

### *Pgbd5* is expressed in embryonic mouse neocortex and is involved in neuron generation

Gene expression analysis from public databases (GTEx for humans and RIKEN FANTOM5 project for mice) indicates that *Pgbd5* is mainly expressed in the brain both in mouse and in human tissues (Fig. 1a). To assess *Pgbd5* expression during embryonic brain development, we performed *in situ* hybridization on coronal brain sections from mouse embryos. As early as E12.5 and consistently at E14.5, *Pgbd5* expression was detected throughout the mouse developing cerebral cortex from the ventricular zone (VZ) to the pial surface (Fig. S1a). To investigate the role of *Pgbd5* during brain development *in-vivo*, we knocked down (KD) *Pgbd5* mRNA by *in utero* electroporation (*iue*) (Fig. 1b). We injected a short hairpin RNA (shRNA)-expressing plasmid targeting mouse *Pgbd5* RNA (shPgbd5) at E12.5, right after the start of cerebral cortex neurogenesis. As a control, we used a non-targeting shRNA (Addgene plasmid #69040, shCtrl), which does not match any known sequences in the mouse or human transcriptomes. Potential off-target effects of both shRNAs were assessed via BLASTN analysis against the mouse transcriptome (Ensembl release 109) ^17^. Only shPgbd5 displayed a perfect 21-nt match exclusively to the *Pgbd5* transcript, ruling out potential off-targets.

**Fig. 1.**
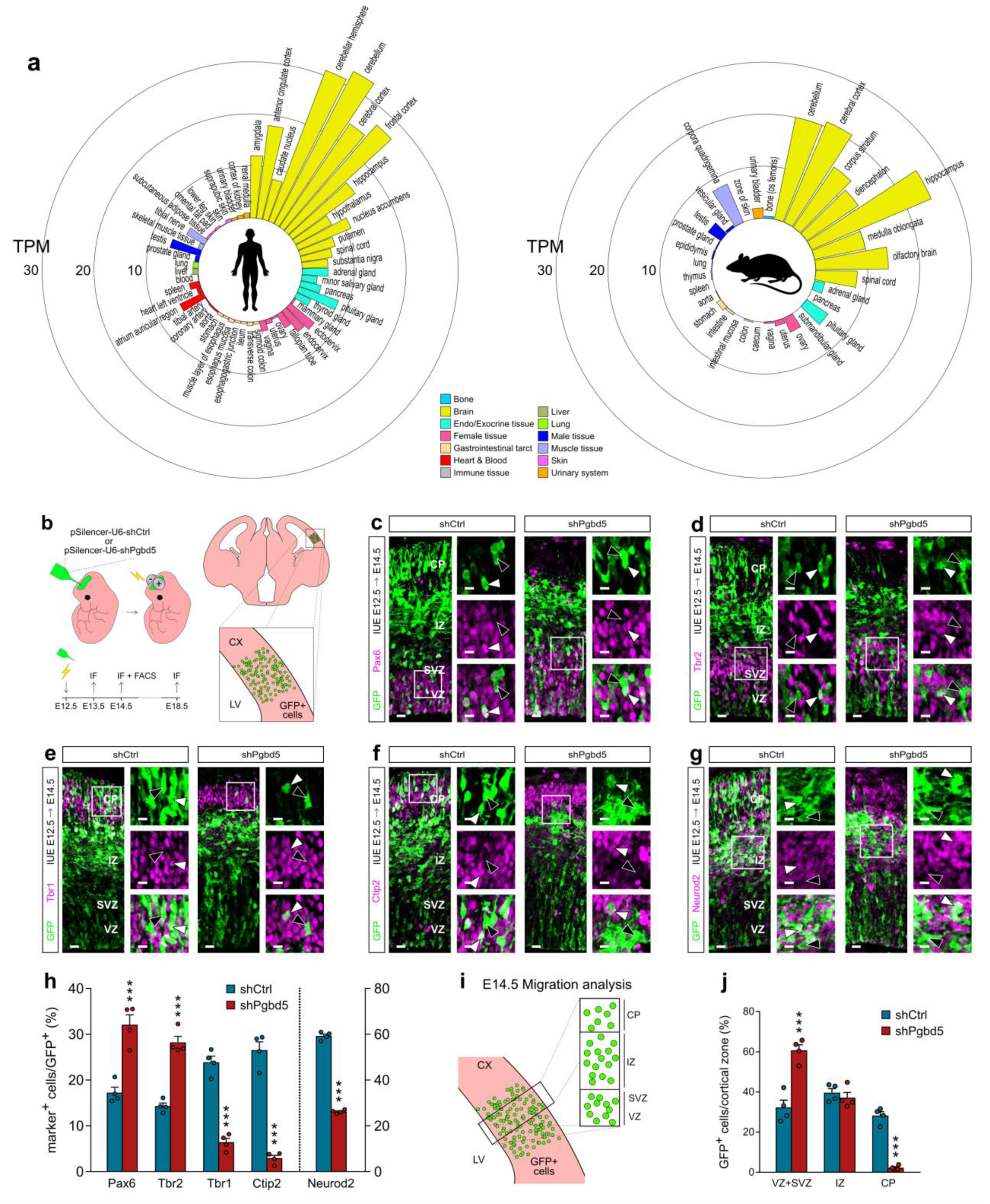
Pgbd5 knockdown alters cellular composition and neuron migration in the cerebral cortex . **a**, Circular barplot of *Pgbd5* expression in human (left) and mouse (right). Expression levels are indicated as transcripts per million reads (TPM). **b**, Schematic of *in utero* electroporation (iue) procedure (top left), experimental timeline (bottom left), electroporated GFP^+^ cells in a coronal brain slice (top right) and electroporated cerebral cortex close-up (bottom right). **c-g**, IF staining for NPCs (Pax6, Tbr2), neuronal cells (Neurod2) and DL neurons (Tbr1, Ctip2) markers at E14.5, shRNAs iue at E12.5. Positive expression (white arrowheads) vs. no expression (black arrowheads). Scale bar 20um, close-up scale bar 10um. **h**, Quantification of marker^+^/GFP^+^ cells at E14.5. Data are mean ± s.e.m.; n = 4 each. Unpaired Student’s t-test. **i**, Schematic of migrationanalysis. **j**, E14.5 GFP^+^ cells distribution in VZ+SVZ, IZ and CP of E12.5 shCtrl- vs. shPgbd5-injected brains. Data are mean ± s.e.m.; n = 4 each. Two-way ANOVA with Sidak’s multiple comparison correction. VZ, ventricular zone; SVZ, subventricular zone; IZ, Intermediate zone; CP, cortical plate; LV, lateral ventricle; CX, cerebral cortex. *P < 0.05, **P < 0.01, ***P < 0.001.

To visualize targeted cells, a vector expressing GFP was co-electroporated at a 1:1 molar ratio. Under these conditions, we achieved a ∼75% reduction of *Pgbd5* RNA expression, as determined by RT-qPCR on FACS-purified GFP^+^ cells at E14.5 (Fig. S1c).

To study the acute effects of *Pgbd5* KD on NPCs and neurons, we collected brains at E13.5, 24 hours post-electroporation. We found a significant decrease in the proportion of GFP^+^ cells expressing neuronal markers (Neurod1 and Neurog2), without significant changes in the number of Pax6^+^ and Tbr2^+^ cells (Fig. S1d-h). Later on, in order to better evaluate the effect of *Pgbd5* KD, we collected brains at E14.5, 48 hours post-iue. At this stage, the cerebral cortex is more expanded and immature neurons are migrating through the IZ and starting to populate the DL of the developing CP. We analyzed GFP^+^ cells by immunofluorescence (IF) using canonical cerebral cortex markers (Fig. 1c-g). Upon *Pgbd5* KD, we observed a significant increase in the proportion of GFP^+^ cells expressing markers of NPCs (Pax6 and Tbr2) compared to shCtrl-injected cortices (Fig. 1h). Conversely, the percentage of GFP^+^ cells expressing the neuronal marker Neurod2, typically found in IZ+CP, was significantly reduced (Fig. 1h). Importantly, staining for cleaved Caspase-3 showed no differences between conditions, ruling out apoptosis as a contributing factor (Fig. S1b). These data suggest that *Pgbd5* expression is required during early cortical development to promote the transition from progenitors to neurons.

In addition, we analyzed early neuronal migration by subdividing the neocortex into three main zones: CP, IZ and VZ+SVZ (Fig. 1i). We quantified the distribution of GFP^+^ cells at E14.5 within these compartments. In *Pgbd5 KD* areas, GFP^+^ cells accumulated predominantly in the VZ+SVZ, and the proportion of migrating neurons reaching the developing CP was significantly reduced compared to controls (Fig. 1j). Furthermore, when analyzing the entire GFP^+^ population, we observed a marked decrease in the number of cells expressing DL markers Tbr1 (L6 marker) and Ctip2 (L5 marker) in *Pgbd5* KD cells compared to shCtrl-treated brains (Fig. 1h). These findings further support an impairment in neuronal positioning upon *Pgbd5* KD.

Together, these results indicate that *Pgbd5* is essential during cortical development for the generation of neurons capable of initiating proper radial migration throughout the neocortical wall.

### *Pgbd5* re-expression can partially rescue neurons generation in *Pgbd5* KD brains

To further confirm the role of *Pgbd5* in cerebral cortex development, we performed rescue experiments at E12.5 by co-electroporating a plasmid encoding a Myc-tagged, shRNA-resistant mouse *Pgbd5* coding sequence (CDS) together with shPgbd5 at a 1:1 molar ratio. Brains were collected at E14.5 and GFP^+^ cells were analyzed by IF, following the same procedure used for shCtrl- and shPgbd5-injected brains (Fig. S2a). On average, ∼75% of GFP^+^ cells expressed Myc-tagged *Pgbd5* (Fig. S2c). Notably, re-expression of *Pgbd5* restored the proportion of GFP^+^ cells expressing Neurod2 and Tbr2 (Fig. S2b), while Pax6, Tbr1 and Ctip2 expressing GFP^+^ cells remained unchanged. Analysis of the spatial distribution of rescued GFP⁺ cells showed that the fraction of *Pgbd5*-expressing cells (MycTag^+^) retained in the VZ/SVZ, reached levels comparable to shCtrl-injected brains. Despite the incomplete restoration of *Pgbd5*-expressing neurons within the CP, the enrichment of rescued cells in the IZ points to a regained migratory competence (Fig. S2d). Notably, the fraction of MycTag⁻ cells remained largely confined to the VZ/SVZ, mirroring the distribution pattern of *Pgbd5* KD cells. This finding highlights the essential role of *Pgbd5* in controlling neuronal positioning during cortical development.

### *Pgbd5* knockdown impairs radial migration of neocortical neurons

To further investigate the impaired cell migration observed at E14.5, we analyzed shPgbd5-injected brains at E18.5, a stage when the layered structure of the cerebral cortex is nearly complete, and the majority of DL neurons have reached their final cortex positions. IF analysis (Fig. 2a) revealed a significant reduction in the number of *Pgbd5* KD GFP^+^ cells expressing the L6 marker Tbr1 compared to controls (Fig. 2b).

**Fig 2.**
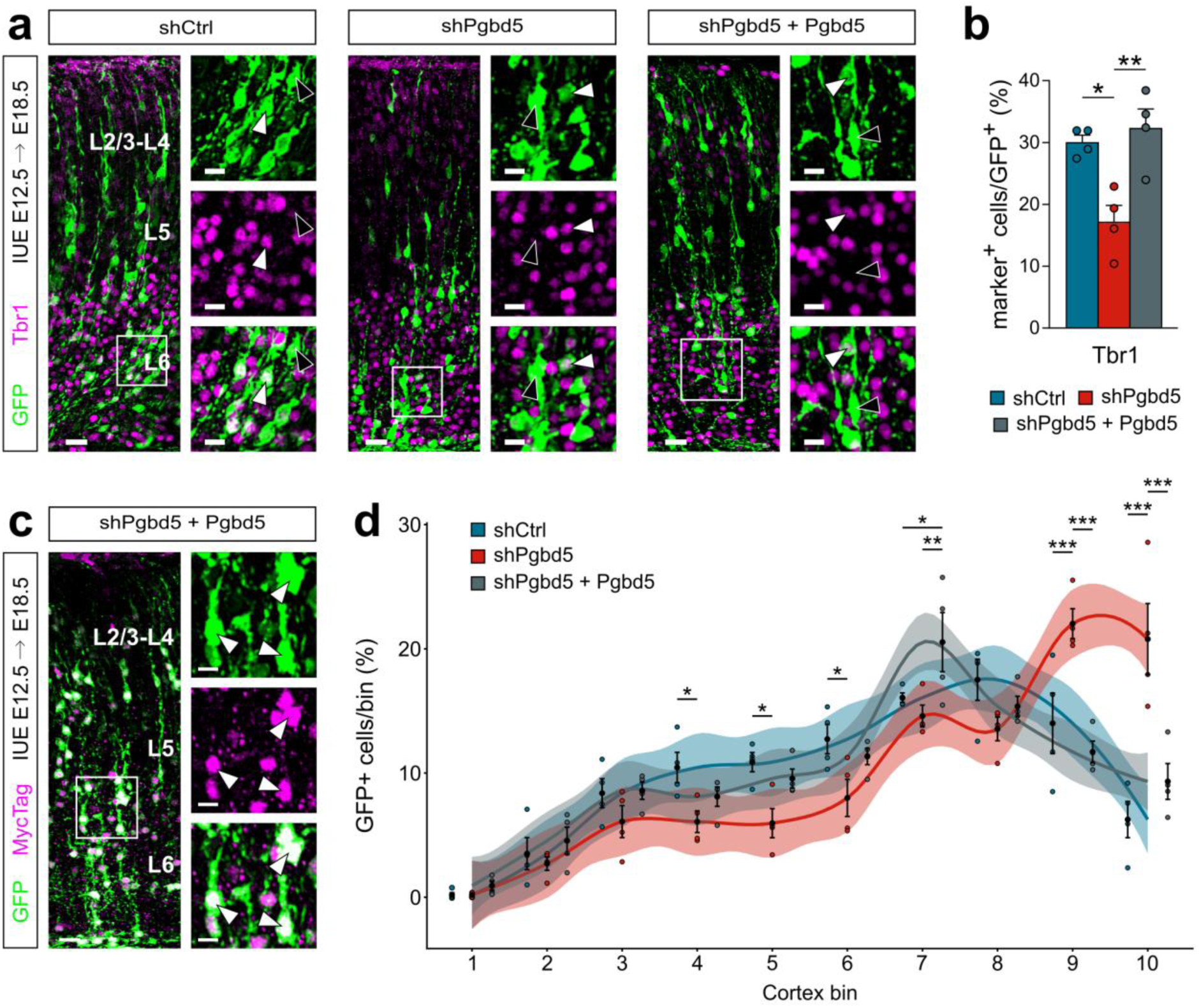
Cerebral cortex neurons migration is rescued after Pgbd5 expression in Pgbd5 KD cells. **a**, IF staining for DL neuron marker Tbr1 at E18.5, shRNAs *iue* at E12.5. Positive expression (white arrowheads) vs. no expression (black arrowheads). Scale bar 20um, close-up scale bar 10um. **b**, Quantification of Tbr1^+^/GFP^+^ cells at E18.5. Data are mean ± s.e.m.; *n* = 4 each. One-way ANOVA with Tukey’s multiple-comparisons test. **c**, IF staining for MycTag^+^ electroporated cells. Scale bar 20um, close-up scale bar 10um. **d**, E18.5 radial distribution of GFP^+^ cells in the cerebral cortex of E12.5 shRNAs-injected embryos. Data are mean ± s.e.m.; *n* = 4 each. Two-way ANOVA with Tukey’s multiple comparison correction. Shaded areas represent the 95% confidence interval for the fitted curve. L6, Layer 6; L5, Layer 5; L2/3-L4, Layers 2/3 and Layer 4; \**P* < 0.05, \*\**P* < 0.01, \*\*\**P* < 0.001.

To further characterize the final positioning of GFP^+^ *Pgbd5* KD cells, we subdivided the cortex into 10 bins and quantified the distribution of GFP^+^ cells in each bin. In *Pgbd5* KD brains, we observed a significant accumulation of GFP^+^ electroporated cells in the lower cortical regions compared with controls (Fig. 2d), indicating that the migration defect detected at earlier stages persisted. Notably, the migration defects observed at E18.5 were partially rescued by co-electroporating Myc-tagged shRNA-resistant mouse *Pgbd5* CDS (Fig.2c). Exogenous *Pgbd5* expression almost completely restored proper cortical positioning of GFP^+^ cells (Fig.2d), as well as the expression of the L6 marker Tbr1 (Fig.2b). Taken together these data suggest that *Pgbd5* is essential for proper neuronal migration through the cortical layers, enabling neurons to reach their final positions within the cerebral cortex.

### *Pgbd5* knockdown impairs cell metabolism and neurogenesis

To identify the molecular pathways affected by *Pgbd5* KD, we performed RNA sequencing (RNA-Seq) on FACS-purified E14.5 GFP^+^ cells from E12.5 shPgbd5- and shCtrl-injected cortices (Fig. 3). *Pgbd5* KD led to the downregulation of 4041 genes and upregulation of 3318 genes (Table S1). Gene Ontology (GO) analysis revealed that upregulated genes were significantly enriched in the regulation of “cell cycle” (e.g. *Ccnd* genes, *Ccng* genes, *Cdk* genes) (Fig. 3a and Table S2). Notably, among the upregulated genes associated with neocortex neurogenesis and patterning, we identified markers of NPCs such as *Pax6*, *Tbr2*, *Nestin*, the Notch pathway genes *Notch1* and *Notch3* and their targets *Hes1* and *Hes5* (Fig. 3c). Remarkably, the upregulation of Notch pathway targets was accompanied by a marked downregulation of proneural genes from the *Neurod* family *(Neurod1*, *Neurod2* and *Neurod6*) consistent with our previous IF findings. Additionally, we observed significant enrichments among downregulated genes in cellular metabolism categories, particularly mitochondrial “oxidative phosphorylation” and “respiratory electron transport chain” (e.g. *Cox* genes, *Nduf* subunits, *Atp5* subunits) as well as “translation” and “ribosome biogenesis” (e.g. *Rpl* and *Mrpl* genes, *Rps* genes) (Fig. 3b and Table S2). Among the significantly downregulated genes, we identified *Ctip2*, *Fezf2*, *Lmo4* and *Satb2*, which are key neocortical layer markers involved in neuronal specification and laminar positioning (Fig. 3c). Furthermore, genes that regulate neuronal migration ( *Dcx*, *Rnd2*, *Rac1*, *Rac3*, *RhoA*, *RhoB*) and cell morphology/polarity (*Sox11* and *Cdc42*) were significantly down-regulated, further corroborating the impaired cell positioning observed at E14.5 (Fig. 3c and Table S2). These findings, at a global transcriptional level, confirm and strengthen our previous observations from IF analysis, further supporting the idea that *Pgbd5* plays a crucial role in regulating cell fate and migration during mouse neocortex development.

**Fig 3.**
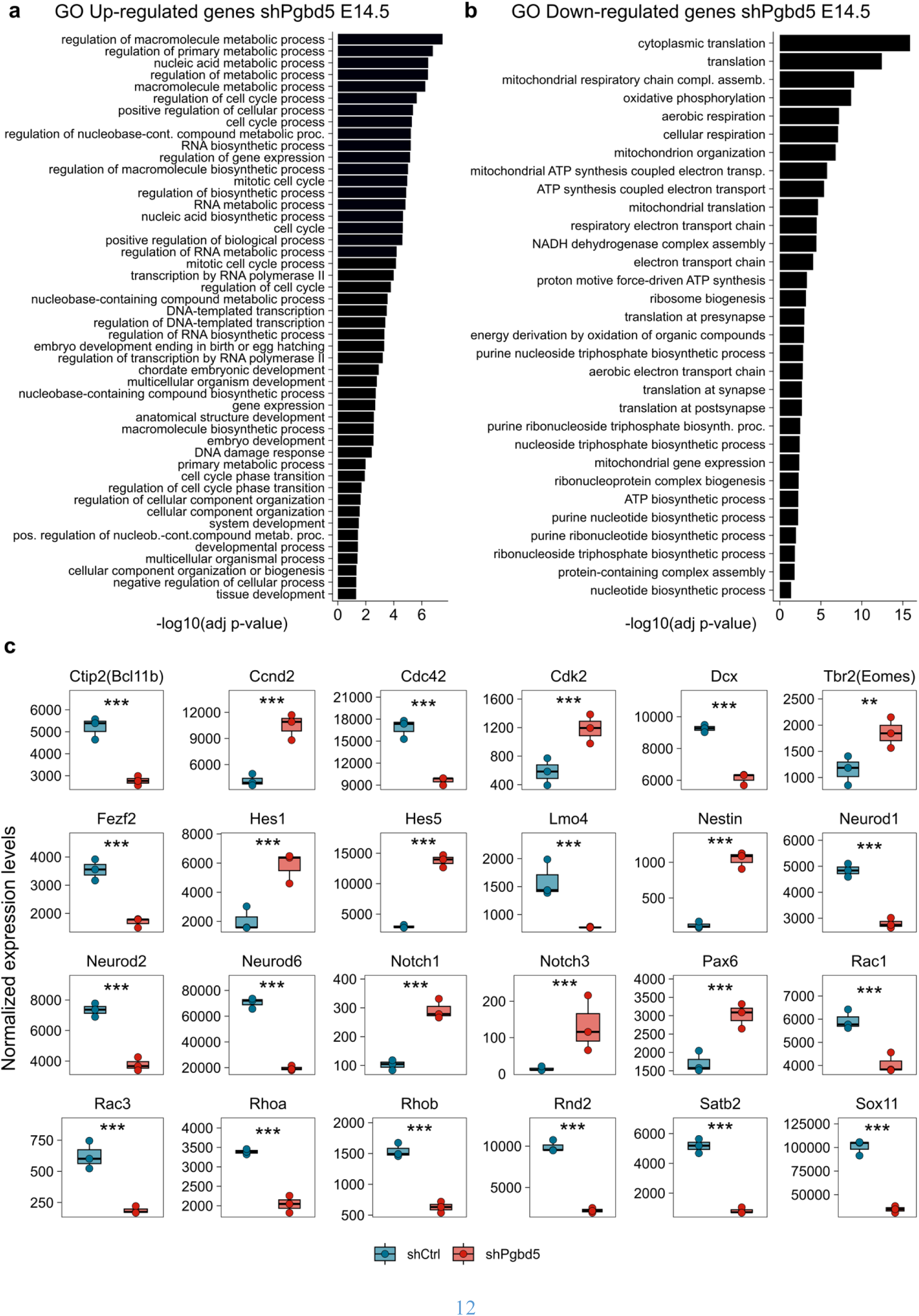
Pgbd5 knockdown impairs cell metabolism and neurogenesis. **a-b**, Significant GO terms (p<0.05) under the biological process category for up-regulated (a) and down-regulated (b) genes in mouse GFP^+^ cells after *iue* of shPgbd5 vs. shCtrl. **c**, Expression values of cerebral cortex gene markers. Boxplots reportingthe normalizedexpressionlevels of specific NPCs and neuronal gene markers. Statistics is reported as FDR-corrected p-value from the DESeq2 DE analysis (see methods). GO, Gene Ontology. \*\**P* < 0.01; \*\*\**P* < 0.001.

### *Pgbd5* knockdown affects physiological occurrence of γH2AX foci in cells

A hallmark of NPCs in the VZ and SVZ is the frequent presence of DSBs ^13,14^. Given that *Pgbd5* possesses endonuclease activity ^7^, we investigated whether *Pgbd5* KD affects the levels of physiological DSBs in NPCs. IF staining of the DNA DSBs marker γH2AX revealed a robust signal in neocortical VZ and SVZ NPCs of wild-type (WT) mice (Fig. 4e). In shCtrl-injected embryos, the γH2AX signal pattern was comparable to WT conditions, ruling out surgery-related artifacts (Fig. 4a). By contrast, γH2AX signal was significantly reduced in the GFP^+^ area of *Pgbd5* KD brains (Fig. 4b). Quantification confirmed a significant decrease in γH2AX^+^ GFP^+^ cells in the VZ and SVZ upon *Pgbd5* KD (Fig. 4d) and γH2AX signal in VZ+SVZ increased after *Pgbd5* re-expression in *Pgbd5* KD cells (Fig. 4c). The dependence of DNA DSBs signal upon *Pgbd5* KD was also further validated *in-vitro* in N2a cell cultures (Fig. S3a). Following *Pgbd5* mRNA KD (Fig. S3b), western blot analysis showed a significant reduction in γH2AX protein levels compared with shCtrl treated cultures (Fig. S3c).

**Fig 4.**
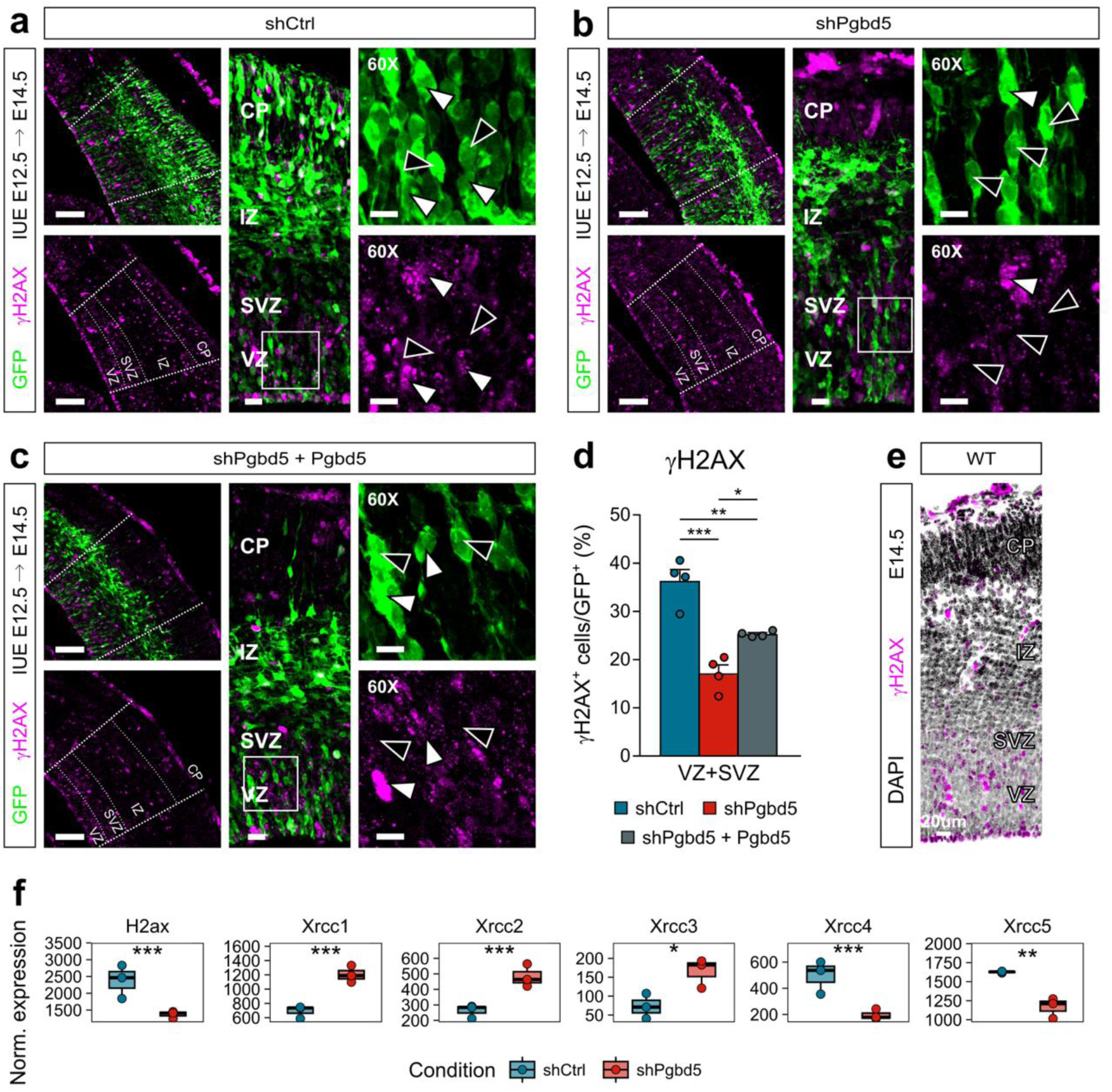
Pgbd5 knockdown affects physiological occurrence of DNA double-strand breaks in cells. **a-c**, γH2AX IF signal in shCtrl- (a) vs. shPgbd5- (b) vs. *Pgbd5* rescue-injected cortices (c). Positive expression (white arrowheads) vs. no expression(blackarrowheads). Scale bar iue area 100um, scale bar 20um, close-upscale bar 10um. **d**, Quantificationof γH2AX^+^/GFP^+^ cells at E14.5. Data are mean ± s.e.m.; *n* = 4 each. One-way ANOVA with Tukey’s multiple-comparisons test. **e**, γH2AX IF staining in WT mouse cerebral cortex at E14.5. **f**, Expression values of DNA damage response gene markers. Boxplots reporting the normalized expression levels. Statistics is reported as FDR-corrected p-value from the DESeq2 DE analysis (see methods). VZ, ventricular zone; SVZ, subventricular zone; IZ, Intermediate zone; CP, cortical plate. \**P* < 0.05, \*\**P* < 0.01, \*\*\**P* < 0.001.

The nucleoside analog 5-ethynyl-2’-deoxyuridine (EdU) is known to trigger DNA damage signaling, such as phosphorylation of histone H2AX on Ser139 (γH2AX), once incorporated into DNA ^18,19^. Consistently, EdU injection 24 hours post-iue triggered a robust increase of γH2AX^+^ staining in control compared to uninjected cortices (Fig. S3d). Interestingly, a significant decrease in the number of γH2AX^+^ GFP^+^ cells was observed following *Pgbd5* KD in the presence of EdU-induced DNA damage signaling ^18,19^ (Fig. S3d–f). Together, these findings highlight a role for *Pgbd5* in affecting DNA DSBs signaling, both in physiological and DNA damage conditions.

By examining our RNA-Seq dataset for genes involved in DNA damage responses, we found that *H2ax*, *Xrcc4* and *Xrcc5*, key components of DSBs repair and of the NHEJ pathway, were significantly downregulated upon *Pgbd5* depletion (Fig. 4f and Table S1). These results are in line with recent work from the Kentsis group linking *Xrcc5* to Pgbd5 activity ^20^. In contrast, genes associated with alternative repair pathways were upregulated in shPgbd5 samples, including *Xrcc1,* primarily involved in base-excision repair (BER), and *Xrcc2* and *Xrcc3*, which participate in homologous recombination (HR) (Fig. 4f and Table S1).

### Loss of *Pgbd5* does not impact somatic genomic variations at E14.5

To explore whether *Pgbd5* induces somatic variants during corticogenesis, we performed ultra deep whole genome sequencing (WGS) on E14.5 GFP^+^ cells sorted from mouse embryos treated with either shCtrl or shPgbd5 at E12.5. Although somatic variants are expected to occur in a small subset of cells, and their detection might be challenging in bulk sequencing, we hypothesized that ultra-deep sequencing could still reveal some of them. We performed PCR-free Illumina WGS with a mean coverage of 170X and subsequently characterized the somatic variant landscape of shCtrl and shPgbd5 treated cells, identifying: i) single nucleotide variants (SNV); ii) insertions and deletions (INDELs); iii) structural variants (SVs) and iv) L1 insertion sites (L1 INS), using computational methods specifically designed for somatic variant detection (see Methods).

We observed that SNVs were the most frequent somatic variants, with a median of 60,000 events per sample, followed by SV (20,000 events/sample) and INDEL (1,000 events/sample) (Fig. 5a). Among the SNVs, G>A and C>T substitutions were most common, while INDELs were predominantly categorized as deletions (DEL, ∼86%) and insertions (INS, ∼13%). SVs were primarily BNDs (complex re-arrangements, ∼78%), deletions (DEL, ∼15%) or duplications (DUP, ∼7%) with very few insertions and LINE-1 insertions (Fig. 5b). Mutational signature analysis revealed an association between somatic SNVs and clock-like SBS5 mutational signature in both *Pgbd5* KD and control conditions (data not shown).

**Fig 5.**
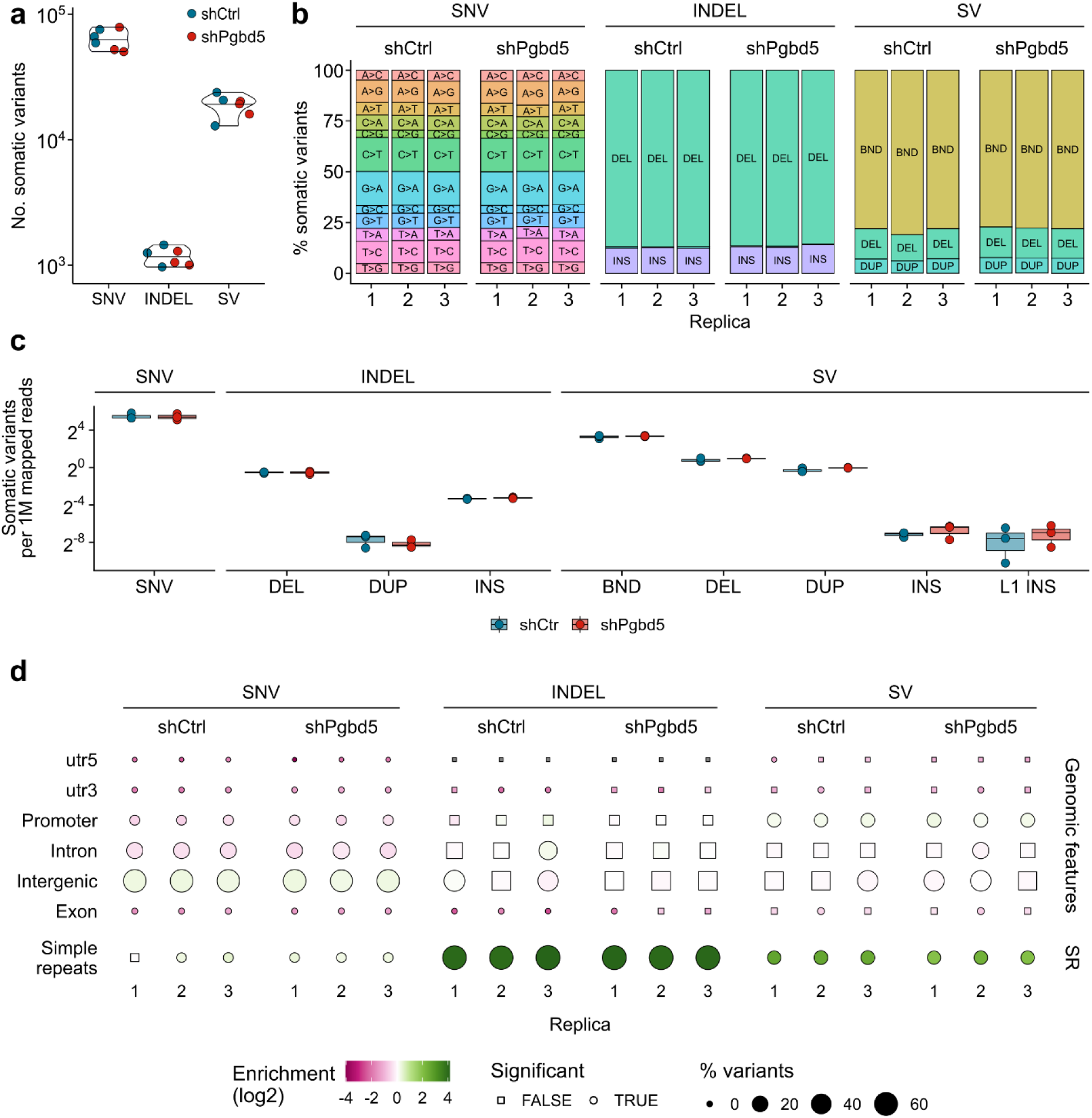
Loss of Pgbd5 has no effects on somatic genomic mutations. **a,** Raw number of somatic variants per sample. The raw number of somatic variants (SNV, INDEL, SV) is reported for each biological replica (n=3) and condition (shCtr and shPgbd5). **b,** Percentage of somatic variants composing each set of event. For each type of variant (SNV, INDEL, SV), the percentage of variants belonging to different variant types is reported. To increase readability of the figure, only variants representing more than 1% of the total variants were labelled. As such, INDEL duplications and SV insertions and L1 insertions are reported in the bars, but not labelled. **c,** Normalized number of somatic variants per sample. The raw number of somatic variants (SNV, INDEL, SV) was normalized on the number of mapped reads, multiplied by 1,000,000 and reported for each biological replica (n=3) for both conditions (shCtrl and shPgbd5). **d,** Enrichment of the somatic variants in genomic features and simple repeats. Each set of variants of each individual replica was overlapped with genomic features (promoter [+/-3 kb from transcript TSS], intron, exon, utr5, utr3, intergenic) and simple repeats. To calculate the enrichment (zscore), the same analysis was repeated on variant coordinates randomly placed in the murine genome 1,000 times each. Zscores were converted to p-value, which were, in turn, FDR-corrected (Benjamini-Hochberg). Fill colours indicate enrichment (zscore) over randomly placed variants. Dot size indicates the percentage of somatic variant overlapping each feature. Dot shape indicates whether the enrichment or depletion over random variants is statistically significant (p<0.05, circle) or not (p>0.05, square).

To compare the somatic events in controls and *Pgbd5* KD conditions, we normalized the number of somatic variants on the sequencing depth of each sample. No statistically significant differences were observed in the normalized number of any type of genomic variation upon *Pgbd5* KD (Fig. 5c).

Next, we sought to determine whether the somatic variants were localized differently in genomic features and simple repeats in *Pgbd5* KD versus control samples. We overlapped the genomic coordinates of each set of somatic variants with the features of interest and repeated the analysis 1,000 time with variants of the same length randomly placed in the reference genome. As expected, somatic SNV were significantly depleted from promoter and intronic region and enriched in intergenic regions (Fig. 5d). This was observed for both controls and *Pgbd5* KD samples. Overall, no significant differences were observed in the localization relative to genomic features of any type of somatic variant between the two conditions (Fig. 5d). Similarly, the location of the somatic variants in annotated simple repeats did not differ significantly between *Pgbd5* KD and control samples across all variant types analyzed (Fig. 5d). Yet, all the types of somatic variants were more enriched in simple repeats than expected by chance, in both control and *Pgbd5* KD conditions, with up to the 75% of the somatic INDELs overlapping annotated simple repeats (Fig. 5d). This same pattern was observed in different murine adult tissues (Ansaloni and Simi *et al.*, under submission), suggesting that somatic expansion and contraction of simple repeats represent an important source of somatic variation.

In conclusion, our results indicate that *Pgbd5* knockdown does not lead to detectable genomic alterations during corticogenesis at E14.5.

## DISCUSSION

Neurogenesis, the process by which dividing NPCs generate post-mitotic neurons, relies on stereotyped and evolutionary conserved molecular pathways. Proper regulation of the balance between proliferation and differentiation is essential for the establishing the layered structure of the cerebral cortex. The cytoarchitecture of the neocortex is tightly orchestrated during embryonic development and further refined postnatally through genetic, epigenetic and activity-dependent mechanisms ^21–23^. In this context, neuronal delamination, migration and positioning are crucial steps, and perturbations in these processes can lead to neurodevelopmental disorders^24^.

Our study identifies *Pgbd5*, the most evolutionarily conserved transposase-derived nuclease gene in vertebrates ^6^, as a key player in cerebral cortex formation.

Acute *Pgbd5* knockdown during the early neocortical development *in vivo* resulted in a cell-autonomous impairment of neurogenesis. Both immunohistochemistry and transcriptomic analysis revealed a marked downregulation of genes involved in neuronal determination and maintenance in the neocortex (e.g. *Neurod* genes, *Dcx*) accompanied by upregulation of proliferation-associated genes (e.g. *Notch* genes, *Hes* genes, *Nestin*, *Cyclin* genes), typical of NPCs ^25^.

Furthermore, *Pgbd5* KD led to migration defects as early as two days post-electroporation, with neurons displaying aberrant positioning within the developing cortex. This was confirmed by migration analysis and RNA-seq, which showed significant downregulation of neocortical migration-controlling genes, such as small *GTPases* ^26^. These defects became more pronounced at perinatal stages, with neurons failing to properly migrate through the cortical plate.

Importantly, re-expression of *Pgbd5* rescued the impaired migration phenotype observed at later developmental stages, underscoring its essential roles in neurogenesis and cortical organization. A recent independent study reported that *PGBD5* knockdown impairs glioma cell migration and invasion *in vitro*, further supporting a role for this gene in regulating cellular dynamics ^27^.

At E14.5, *Pgbd5*-mediated rescue was incomplete, with a partial restoration of the proportion of cells reaching the CP. This partial rescue may reflect the concentration of the plasmid used for rescue and/or the differing turnover rates of Pgbd5 and GFP proteins. Considering that approximately 75% of GFP⁺ cells were expressing MycTag, the significant increase in MycTag⁺ GFP⁺ cells within the IZ, together with the restored expression of *Neurod2*, indicates that *Pgbd5*-expressing cells are engaging in appropriate migratory processes, potentially with a slight delay, and are likely to reach their correct cortical destinations by perinatal stages.

GO analysis of the differentially expressed genes revealed a strong impact of *Pgbd5* downregulation on cellular metabolism. Numerous genes encoding subunits of the mitochondrial oxidative phosphorylation (OXPHOS) complexes ^28^ were significantly decreased following *Pgbd5* knockdown. During the transition from NPCs to neurons, cells progressively shift from glycolysis to OXPHOS to meet the energy demands of neuronal differentiation and function rather than proliferation ^29–31^. It remains to be determined whether this metabolic switch is directly controlled by *Pgbd5* or represents a secondary consequence of an altered balance between proliferation and differentiation.

*Pgbd5* knockdown, both *in vitro* and *in vivo*, results in a marked reduction in γH2AX foci, a well-established marker of DNA DSBs formation and repair. These findings suggest that *Pgbd5* contributes directly or indirectly to a physiological pathway involving DSBs during corticogenesis. This interpretation is supported by independent evidence from the Kentsis’s group ^20^, who similarly observed reduced γH2AX foci in the developing neocortex of *Pgbd5* knockout mice and demonstrated that DSB formation requires a functional endonuclease domain. As a note of caution, we cannot exclude a more conservative interpretation of our results where the reduction in γH2AX foci may simply reflect the downregulation of H2AX itself and therefore of γH2AX immunodetection upon *Pgbd5* depletion.

Ultra-deep *in vivo* WGS did not reveal significant quantitative or qualitative differences in somatic variant profiles between *Pgbd5*-KD and control samples at E14.5. This outcome may reflects the intrinsically low frequency of somatic events and the associated limitations of detecting such variants by bulk Illumina WGS. It is also plausible that the *Pgbd5*-induced DSBs detected by immunofluorescence are rapidly and accurately repaired, leaving no persistent genomic alterations identifiable by short-read sequencing and variant-calling pipelines. Although more specific and *ad hoc* investigations will be required, a recent study examining the resolution of Cas9-induced double-strand breaks *in vitro* points to a similar mechanistic direction, at least with respect to DSB repair in post-mitotic neurons. In this work, the authors report that post-mitotic neurons either require substantially longer time to resolve DSBs or undergo a more accurate and cleaner repair process compared to proliferating cells ^32^.

DSBs are mainly repaired by HR, an error-free mechanism active in proliferating cells, and NHEJ, which operates in both neural progenitors and differentiated post-mitotic neurons and may introduce small errors. When HR and NHEJ are insufficient, cells engage additional, more error-prone repair pathways ^33,34^. Canonical NHEJ is also considered the principal mechanism resolving DSBs generated by PiggyBac-like transposases ^35–37^.

In shPgbd5 cells, *H2ax*, *Xrcc4* and *Xrcc5*, key components of canonical NHEJ, are downregulated, consistent with a reduced basal capacity for DSB repair. H2AX, a conserved H2A variant, is rapidly phosphorylated (γH2ax) during early repair. XRCC4 stabilizes and recruits Lig4 ^38^ to DSBs ^39,40^, whereas *Xrcc5* encodes Ku80, a core subunit of the Ku heterodimer that binds DNA ends to initiate NHEJ ^41^. Consistently, Kentsis and colleagues recently showed that *Xrcc5* is required for repairing *Pgbd5*-dependent DSBs ^20^.

Conversely, induction of *Xrcc1*, *Xrcc2* and *Xrcc3* suggests compensatory engagement of alternative repair pathways. It is also possible that DSBs generated between E12.5 and E14.5 persist beyond the sampling window and thus escape detection.

Although DNA-damage responses are predominantly regulated at the post-translational level, sustained transcription of core NHEJ components may represent an additional control layer in long-lived neurons. In this context, Pgbd5 may both facilitate physiological DSB formation ^42^, and modulate the machinery required for their repair.

The role of *Pgbd5* in corticogenesis aligns with broader evolutionary hypotheses in which transposon-derived genes, originally selfish elements, become co-opted and subfunctionalized to perform beneficial host functions. Enhancing the robustness of DSB repair could confer advantages both to the host genome, by supporting the long-term integrity of neurons, and to PiggyBac-derived machinery, whose life cycle depends on accurate resolution of transposition-associated DNA breaks.

## METHODS

### Gene expression analysis from publicly available data

Expression data from normal tissues was obtained from the GTEx portal v8 for humans and RIKEN FANTOM5 project for mice.

### Animals

CD1 IGS mice (Charles River) for in utero electroporation experiments (*iue*) were housed at Istituto Italiano di Tecnologia (IIT) animal facility. All animal procedures were approved by IIT animal use committee and the Italian Ministry of Health and conducted in accordance with the Guide for the Care and Use of Laboratory Animals of the European Community Council Directives. All mice were group-housed under a 12-hours light-dark cycle in a temperature and humidity-controlled environment with *ad libitum* access to food and water.

### Constructs

For *iue* experiments, control shRNA (shCtrl) and *Pgbd5* shRNA (shPgbd5) (Table S5) were cloned along the hU6 promoter in a pSilencer 3.0-H1 (ThermoFisher Scientific) plasmid using EcoRI and HindIII restriction sites and substituting the original H1 promoter. To visualize electroporated cells, pSilencer 3.0-U6-shCtrl and pSilencer 3.0-U6-shPgbd5 vectors were co-electroporated together with a GFP expressing pCAGIG vector (Addgene #11159) in a 1:1 molar ratio. For *in-vivo* rescue experiments a NLS- and Myc-tagged shRNA-resistant version of mouse *Pgbd5* was cloned in a pCAGIG vector (GenScript).

### In Utero Electroporation

Animal care and experimental procedures were performed in accordance with the IIT licensing and the Italian Ministry of Health license n° 693/2019-PR and 327/2021-PR. Briefly, E12.5 timed-pregnant CD1 IGS mice were anesthetized with isoflurane and administered 2-5 mg/kg ketoprofene for analgesia. The uterine horns were exposed by laparotomy and the DNA plasmids with 0.01% Fast Green dye (Sigma Aldrich) were injected in the lateral ventricle using a glass capillary (B100-58-10, Sutter Instrument). 3 mm diameter platinum tweezer electrodes (CUY650P3, NepaGene) were used to electroporate the cerebral cortex. Four electrical pulses (35 V, 50 msec duration, 950 msec interval) were delivered using a NEPA21 electroporator (NEPA21, NepaGene). The nucleoside analog EdU (5-ethynyl 20-deoxyuridine) was injected intraperitoneally at 25 mg/kg in pregnant females at E13.5, 24h after the *iue* procedure. EdU was detected using EdU click-iT technology (Invitrogen, C10340).

### Cell cultures

N2a cells (American Type Culture Collection) were maintained in Dulbecco’s modified Eagle’s medium (DMEM, Thermo Fisher Scientific) supplemented with 10% fetal bovine serum (FBS, Thermo Fisher Scientific) and 1% antibiotics (penicillin/streptomycin, Thermo Fisher Scientific) at 37°C and 5% CO2. N2a cells were plated in 6-well plates and transfected with 1ug of 3.0-U6-shCtrl or pSilencer 3.0-U6-shPgbd5 plasmids using Lipofectamine 2000 (Thermo Fisher Scientific) following manufacturer’s instructions. Cells were harvested 48 h after transfections and RNA and proteins were obtained from the same transfection in each biological replicate.

### Western blot

Cells were lysed in radioimmunoprecipitation assay (RIPA) buffer with the addition of protease inhibitor cocktail (Sigma-Aldrich), briefly sonicated, and boiled with 1X Laemmli buffer for 5 min at 95°C. 20 μg of total lysate were resolved by 10% SDS-PAGE TGX pre-cast gels (Bio-Rad) and transferred to nitrocellulose membrane using Trans-Blot Turbo Transfer System (Bio-Rad). Membranes were blocked with 5% non-fat dry milk in TBS/0.1% Tween 20 and incubated with the following primary antibodies: anti-β-actin-HRP 1:10,000 (Sigma-Aldrich) and anti-gH2AX 1:1,000 (Cell Signaling Technology). Proteins of interest were visualized with the SuperSignal West Pico PLUS Chemiluminescent Substrate (Thermo Fisher Scientific). Western blotting images were acquired with ChemiDoc MP Imaging System (Bio-Rad), and band intensity was calculated using ImageJ software.

### Immunofluorescence and histology

Mice were sacrificed at indicated ages and the brains dissected and fixed in 4% paraformaldehyde (PFA) (Sigma Aldrich) at 4°C overnight. Brains were de-hydrated in 30% sucrose and sliced in 20 µm coronal cryosections using a CM 3050S cryostat (Leica). Sections were stored at −80°C. Slices were permeabilized and blocked in 1x Phosphate-Buffered Saline (PBS) containing 5% normal goat serum (NGS, Abcam) and 0.3% Triton X-100 (Sigma Aldrich). Brain sections were incubated overnight with primary antibodies (Table S3) diluted in blocking solution at 4°C. After extensive washes in 1x PBS + 0.1% Triton X-100 (PBST) brain slices were incubated with fluorescent dye conjugated secondary antibodies (Table S3) diluted in blocking solution for 1h at RT. Slices were counterstained with Hoechst 33342 (Thermo Fisher Scientific) 1:10.000 in PBST for 15 min and extensively washed in 1x PBS, mounted with Mowiol (Sigma Aldrich).

### RNA In situ hybridization (RNA-ISH)

For RNA *in situ* hybridization, RNAScope probes specific for *Pgbd5* were used (ACD, # 561941). Mice were sacrificed at indicated embryonic stages and the brains dissected and fixed in 4% paraformaldehyde (PFA) (Sigma Aldrich) at 4°C overnight. Brains were de-hydrated in 30% sucrose and sliced in 12 µm coronal cryosections using a CM 3050S cryostat (Leica).

Sections were stored at −80°C. The RNA-ISH was performed following manufacturer’s instructions. Briefly, cryosections were baked for 30 min at 60°C and fixed in 4% PFA for 15 min at 4°C followed by PBS wash. After dehydration, cryosections were treated with hydrogen peroxide for 10 min at RT, target retrieval reagents at 95°C for 15 min and Protease III treatment for 30 min at 40°C. The probe set was hybridized for two hours at 40°C. Signal amplification steps were performed according to the manufacturer’s protocol using TSA Vivid fluorophores (Tocris).

### Imaging and analysis

Fluorescent images were acquired with Nikon A1 confocal microscope equipped with a 20x objective and processed with Nikon software version 4.11.0 (NIS Elements). Positive cells for the indicated marker were counted through the depth of the cortex in the electroporated area and the percentages normalized on the total number of GFP^+^ cells as indicated in the figure legends. For cell number quantifications, all relevant sections containing GFP^+^ electroporated cells from rostral to caudal were quantified upon shCtrl and shPgbd5 conditions by ImageJ version 1.53q (Wayne Rasband, National Institutes of Health, USA). In order to assess GFP^+^ cells distribution at E14.5, the electroporated cortex was subdivided radially in three bins using Tbr1 and Tbr2 markers to identify the CP, IZ or VZ+SVZ boundaries. For the E18.5 migration analysis, we divided the cerebral cortex in ten equal-sized bins. In both analyses the number of GFP+ cells in each bin were quantified and reported as the percentage of total counted cells.

Statistical analyses on IF experiments were performed using GraphPad Prism (version 7, GraphPad Software) according to the number of replicates and group design. In all figures, significance of comparisons is denoted as follows: \**P*<0.05, \*\**P*<0.01, \*\*\**P*<0.001. In the figure legends, statistical tests used and the nature and numbers of samples analyzed (defined as *n*) are reported. Sample sizes were based on published experiments and previous experience in which differences were observed. No power calculations or statistical tests to pre-determine the sample size were used.

### FAC-sorting

For *in-vivo* RNA-seq pregnant female CD1 IGS mice were electroporated at E12.5 with shCtrl or shPgbd5, sacrificed at E14.5 and brain cortices dissected under a stereomicroscope in ice-cold Hanks’ Balanced Salt Solution (HBSS) and enzymatically dissociated using a Neural Tissue Dissociation Kit (Miltenyi Biotec) following manufacturer’s protocol. After tissue digestion at 37°C, cells were manually dissociated by pipetting, filtered with a 40µm cell strainer, centrifuged for 10 min at 300 rcf and resuspended in ice-cold HBSS.

### RNA isolation, RT-qPCR and library preparation

Total RNA was isolated from cells by TRIzol Reagent (Thermo Fisher Scientific) or RNeasy Mini Kit (Qiagen). DNA was removed by treatment with DNAse I (Sigma-Aldrich). cDNA was synthesized from 0.5 ug of RNA using iScript cDNA Synthesis kit (Bio-Rad). Real time quantitative PCR was performed on a CFX96 Touch™ Real-Time PCR Detection System (Bio-Rad). For *Pgbd5* detection a SsoAdvanced Universal SYBR Green Supermix (Bio-Rad) was used with *Gapdh* as reference gene with the following cycling conditions: 95°C for 20 sec, followed 40 cycles of 95°C for 10 sec and 60°C for 1 min. No template and no RT controls were included. Expression levels were determined relative to the reference gene using the ΔΔCt method.

For *in-vivo* RNA-Seq in each shCtrl and shPgbd5 sample, *n* = 3 pools of 250 GFP^+^ cells were collected for direct RNA-Seq library preparation using the SMART-Seq HT PLUS kit (Takara), according to manufacturer’s instructions.

### Gene expression analysis

RNA-Seq reads were mapped to the murine reference genome (version: GRCm39.vM28) using STAR with default parameters (version: 2.6.0a) ^43^. Gene expression levels were estimated by the read counting module embedded within the STAR tool (–quantMode GeneCounts). DE-genes were identified by using DESeq2 (v.1.48.2) ^44^. Lowly expressed genes were removed, retaining only genes with more than 10 sequencing reads assigned in at least 2, out-of-the 6, sequenced samples. Genes were considered as DE when showing an FDR < 0.05.

### Gene ontology enrichment analysis

Gene ontology (GO) enrichment analysis was performed by using gprofiler2 ^45^. GO terms enriched in the set of up- and down-regulated genes were identified separately providing a custom list of background genes (*i.e.*, expressed genes >=10 raw reads in at least two samples). gSCS p-value correction method was used. GO terms associate to at least 5 up- or down-regulated genes and showing FDR < 0.05 were considered as significant.

### Whole genome sequencing

#### Library Preparation

DNA was quantified with a Qubit fluorometer (Thermo Fisher Scientific, USA). A total of 200 ng of high-quality genomic DNA per sample was used for library preparation. Whole-genome sequencing PCR-free libraries were prepared using the Illumina DNA PCR-Free Prep kit (Illumina Inc., San Diego, CA, USA) following the manufacturer’s protocol. Briefly, genomic DNA was enzymatically fragmented and subjected to end repair and A-tailing, followed by ligation of Illumina’s dual-index adapters (Illumina DNA/RNA UD Indexes v3). Libraries were then purified to remove small fragments and adapter dimers. The final libraries were quantified using a Qubit fluorometer by the single-stranded DNA quantification Qubit™ ssDNA Assay Kit.

### Sequencing

The normalized libraries were pooled equimolarly at a final concentration of 1.7nM and sequenced on Illumina NovaSeq 6000 platform (Illumina Inc., USA) using paired-end 150 bp reads. Sequencing was performed to achieve an average coverage depth of 200X per genome.

### Data Processing and Quality Control

Raw sequencing data were processed using Illumina’s bcl2fastq (v2.20.0) software for base calling and demultiplexing, generating FASTQ files for downstream analysis. Quality assessment of raw reads was performed using FastQC (0.12.1).

Given the high concordance between the two technical replicates of the samples SHCTRL_REP_3 and SHPGBD5_REP_2, only one technical replica was retained discarding SHCTRL_REP_3.2 and SHPGBD5_REP_2.2 from further analyses.

### Annotation files used for WGS analyses

The CD1 mouse strain was used for all the experiments of this study. Since the identification of genomic variants might be affected by the mapping of sequencing reads from one strain to a reference genome of a different strain, we used strain-specific CD1 reference genome and annotation files (gtf and repeatmasker annotation files) from a previous publication ^46^. In addition, a gff file was generated by converting gtf to gff using gffread ^47^.

### TE annotation file

The repeatmasker annotation file was parsed to select only actual TEs, discarding other types of repeats. Only repeats annotated as DNA, LINE, LTR, RC, Retroposon or SINE were retained.

### Simple repeat annotation file

To generate a comprehensive annotation of simple repeats in the CD1 genome, we extracted the items annotated as “Simple_repeat” from the repeatmasker annotation file. In addition, to refine the list of simple repeats, we run Tandem Repeat Finder (trf) ^48^ on the CD1 reference genome (*trf $genome 2 5 7 80 10 50 2000 -l 10 -h -d -ngs*). The output file was then converted to bed format using custom code.

### Reads mapping

Reads were mapped to the CD1 reference genome by using the *fq2bam* tool from NVIDIA Clara Parabricks package ^49^ in a singularity (v1.1.8) container (parabricks-4.0.0-1.simg) with default parameters (*pbrun fq2bam --in-fq $fq1 $fq2 --out-bam $outbam --ref $genome --num-gpus 2*). *fq2bam* wraps *bwa mem* ^50^ and *gatk MarkDuplicates* ^51^ hence first mapping the reads to the reference genome and then flagging the duplicated reads. Duplicated reads were then discarded by *samtools view* (*samtools view -F 1028 -b -o ${outbam}_noDupl.bam $bam*) ^52^.

### Variant calling – SNV

#### Somatic calling

*mutectcaller*, the Clara Parabricks accelerated version of *mutect2* ^53^, was run in somatic single mode (*i.e.*, tumor mode with no matched control sample) on each individual bam file (*pbrun mutectcaller --ref $genome --tumor-name sample --max-mnp-distance 0 --in-tumor-bam $bam --out-vcf ${name}.vcf*). The generated vcf files were then compressed (*bgzip ${name}.vcf*) and indexed (*tabix -p vcf ${name}.vcf.gz*).

#### Panel of normal generation

Having identified the somatic SNV in each sample, a panel of normal (PON) was generated for control and *Pgbd5* KD samples, separately. To this end, first, a *pon_db* was generated (*gatk GenomicsDBImport -R $genome --genomicsdb-workspace-path pon_db $list_of_vcf*). Second, the somatic SNVs were combined into a PON (*gatk CreateSomaticPanelOfNormals -R $genome -V gendb://pon_db -O pon.vcf.gz*) and, third, the PON was indexed (*pbrun prepon --in-pon-file pon.vcf.gz*). This was done separately for control and *Pgbd5* KD samples thus generating one PON for controls and one PON for *Pgbd5* KD.

#### SNV filtering

Having generated one PON for controls and one for *Pgbd5* KD samples, *mutectcaller* was re-run flagging the somatic SNV identified in the PON (*pbrun mutectcaller --ref $genome --tumor-name sample --max-mnp-distance 0 --in-tumor-bam $bam --out-vcf ${name}.vcf --pon pon.vcf.gz*). PON generated from controls was used when running the analysis on *Pgbd5* KD and PON generated from *Pgbd5* KD was used when analysing the control samples. Somatic SNV found in PON were then discarded by running *postpon* (*pbrun postpon --in-vcf ${name}.vcf --in-pon-file pon.vcf.gz --out-vcf ${name}.anno.vcf*). Finally, somatic SNV were additionally refined by i) flagging (*gatk FilterMutectCalls -R $genome -V ${name}.anno.vcf -O ${name}.anno_filter.vcf*) and, ii) excluding false positive somatic SNV with custom code by selecting only the SNV flagged as “PASS”. Only 1-bp-long SNV were retained, discarding all the SNV with a reference or an alternative allele longer than 1 bp. This was done in order not to generate a SNV-INDEL spurious set of SNV. Finally, the set of somatic SNV was additionally filtered by retaining only the SNV scoring a minor allele frequency (MAF) below 5% in all the samples where the SNV was found.

### Variant calling – INDEL and SV

#### Somatic calling

Somatic INDEL and SV were identified by running *manta* in somatic mode ^54^. *Manta* was run with default parameters (*configManta.py --tumorBam $bam --referenceFasta $fasta –runDir ${wfdir}*) except for the minimum variant length tolerated that was set to 15 bp (*minScoredVariantSize = 15*, in the *configManta.py.ini*).

#### Variants filtering

Having identified putative somatic INDEL and SV, *manta* was additionally run in germline mode to identify germline variants (*configManta.py --bam $bam --referenceFasta $fasta --runDir ${wfdir}*). Somatic variants identified also by the germline *manta* analysis were removed from further analyses. Only variants flagged as “PASS” were selected. Variants longer than 10 Mbp were additionally removed.

#### INDEL and SV identification

Variants identified by *manta* were then subdivided in INDEL and SV based on their length. Variants longer than 50 bp, or classified as BND, were classified as SV; variants shorter than 50 bp as INDEL.

#### INDEL parsing

Variants classified as INDEL were next combined across the different samples and refined in order to select a real set of somatic INDEL. INDEL were combined across different samples if sharing the same chromosome, start coordinate and variant type. Combined INDEL were considered as true somatic only when minor allele frequency (MAF) was below 5% in all the samples where the INDEL was found. Finally, INDEL found in both controls and *Pgbd5* KD were discarded – mimicking the PON approach used by mutect2.

#### SV parsing

The same approach used for INDEL was applied to SV as well. SV were combined, only SV showing MAF<5% in all the samples where the SV was found and SV found in both controls and *Pgbd5* KD were discarded. Of note, BND have, by definition, two breakpoints. The way the two breakpoints are reported in the vcf file is likely to be random. As such, the same BND can be sometime be reported as breakpoint1→breakpoint2 and other times breakpoint2→ breakpoint1. To avoid considering these events as different, each BND breakpoint was ordered alphabetically in order to have the same BND always reported with the same breakpoint order.

### L1 insertions

#### Input data

To identify somatic L1 insertions sites, xTea ^55^ was used. However, xTea is not designed to work on species other than human. As such, custom implementation was needed to make it compatible with murine data. First, murine L1 consensus sequences (*A_I*, *Gf_I*, *Tf_I, Tf_II*, *Tf_III*) were retrieved from a previous publication ^56^ and then formatted following the “xTea repeat library preparation” section from the xTea GitHub repository. ((https://github.com/parklab/xTea/tree/master/xtea/rep_lib_prep). Briefly, each L1 consensus was indexed by *bwa index* ^50^, the genomic location of the L1 of interest were extracted from the CD1 repeatmasker file retrieved from a previous publication ^46^ extracting only putative full-length copies (length > 5.5 kbp). Finally, xTea was run on the annotation data generated for each L1 to generate properly formatted input data (*x_TEA_main.py -P -K -p ./ -r $genome -a ${te_name}_rmsk_FL.out -o ${te_name}_rmsk_FL_with_flank.fa -e 100*).

#### Germline calling

xTea was initially run in germline mode to generate a list of L1 germline insertion sites that were used as blacklist during the somatic call of L1 insertion site (*xtea -i $sample -b $bam -x null -p ${outdir}/${l1} -o submit_jobs_${l1}.sh -l ${consensus_dir}/${l1} -r $genome -g $gff --xtea/path/to/xTea/xtea/ -f 1555 -y 32 --lsf -t 12:00:00 -q workq -n 20 -m 80*).

#### Somatic calling

xTea was run in somatic mode on each sample, for each L1 of interest, by providing the coordinates of L1 germline insertion sites as blacklist (*xtea -M -U --nclip 2 --cr 0 --nd 1 --nfclip 1 --nfdisc 1 -i $sample -b $bam -x null -p ${outdir}/${l1} -o submit_jobs_${l1}.sh -l ${consensus_dir}/${l1} -r $genome -g $gff --blacklist $blist --xtea /path/to/xTea/xtea/ -f 1555 -y 32 --lsf -t 12:00:00 -q workq -n 20 -m 80*). Only L1 insertions flagged as “PASS” were retained for further analyses.

#### L1 insertion parsing

Since xTea was run separately on 5 different L1 consensus that are, however, known to share a good sequence homology in specific portions of their sequence, the same insertion sites might have been associated to multiple L1 consensus. To avoid such a redundancy, L1 insertions associated to different L1 consensus were aggregated when sharing the same insertion site coordinates. Consequently, no distinctions were made between L1 insertions of *A_I*, *Gf_I*, *Tf_I, Tf_II* or *Tf_III* elements, but all of them were classified as L1 insertions. Next, L1 insertions were combined across different samples and refined to select a real set of somatic L1 insertions. To this end, L1 insertions were combined across different samples if sharing the same chromosome end and start coordinates. Combined L1 insertions were considered as true somatic when: i) xTea detected the presence of the polyA in the L1 insertion in at least 1 sample, ii) xTea detected the presence of the target site duplication (TSD) in the L1 insertion in at least 1 sample and iii) L1 insertion had a MAF < 5% in all the samples where the L1 insertion was found. Finally, L1 insertion found in both controls and *Pgbd5* KD were discarded – mimicking the PON approach used by mutect2.

### Mutational signature analysis

Mutational signature analysis was run on the entire set of somatic SNV using *signatureanalyzer* ^57^ (*signatureanalyzer -n 50 -t maf -o ./ --reference cosmic3 --hg_build $genome --random_seed 1103 ${name}_snv.maf*).

### Overlap with genomic features

The final sets of somatic SNV, INDEL, SV and L1 insertions were annotated compared to the CD1 genomic features. To this end, first, a txdb object was generated from the CD1 gtf file (*GenomicFeatures::makeTxDbFromGFF*) ^58^. Second, genomic ranges of each genomic feature were extracted from the txdb object (*transcripts, promoters, exons, fiveUTRsByTranscript, threeUTRsByTranscript, intronsByTranscript* functions from the *GenomicFeatures* library).

Promoters were defined as +/- 3kb from each transcript TSS and downstream regions as 300 bp downstream to TTS. Then, by using custom code, each somatic variant was overlapped with promoter, 5’ UTR, 3’ UTR, exon, intron, downstream – in this exact order. Variants that did not overlap any of the above, were annotated as intergenic. To define whether somatic variants were significantly enrichment in specific genomic features, the same analysis was repeated 1,000 times on somatic variants of the same exact length randomly placed in the CD1 genome. To this end, the variant coordinates were first converted to GRanges (r*egioneR::toGRanges*) ^59^, shuffled (*ChIPSeeker::shuffle*) ^60^ and then overlapped with genomic features as previously described.

Having counted the number of somatic variants overlapping each genomic feature, in each sample, for both real and shuffled variants, zscore was calculated [ (number of real variants overlapping the feature of interest – mean number of shuffled variants overlapping the feature of interest) / standard deviation of the number of shuffled variants overlapping the feature of interest]. Zscores were then converted to p-values (*pnorm()*) and then adjusted for multiple testing (*fdr()*).

### Overlap with annotated simple repeats

The final sets of somatic SNV, INDEL, SV and L1 insertions were overlapped with simple repeats. Two annotation files were used: simple repeats extracted from the repeatmasker output and simple repeats annotated *de novo* in the CD1 genome (see above “Simple repeat annotation file”). Since this analysis aimed to define whether the somatic variants overlapped, or not, with simple repeats, the two simple repeats annotation files were simply merged by *bedtools* (*cat $file1 $file2 | bedtools sort -i stdin | bedtools merge -i stdin*). Somatic variants were overlapped with simple repeat coordinates by using *bedtools* and considering a variant as overlapping simple repeats if at least 50% of the length of the variant overlapped at least one simple repeat (*bedtools intersect -f 0.5 -c -a $variants -b $simple_rep*). The same analysis was repeated 1,000 on variants randomly placed in the CD1 genome. Random analysis, zcore, p-value and fdr calculations were performed as previously described.

## RESOURCE AVAILABILITY

### Lead contact

Further information and requests for resources and reagents should be directed to and will be fulfilled by lead contact Stefano Gustincich.

### Materials availability

Any requests for resources and reagents should be directed to lead contact

### Data and code availability

All data and materials supporting the findings of this study are available in the main text or the Supplementary Information, and from the corresponding authors upon request. The raw RNA-Seq and WGS-seq data have been deposited at ENA (European Nucleotide Archive) under the series accession codes XXXXX and XXXX. Any additional information required to reanalyze the data reported in this work paper is available from the lead contact upon request.

## ACKNOWLEDGMENTS

We sincerely thank all members of the SG and RS laboratories and the RNA flagship @IIT for their insightful discussions. We are also grateful to the technical and administrative staff of IIT (especially Eva Ferri, Rosa Maria Cossu, Fabrizio Torri, and Alessandra Sanna). We are indebted to the IIT Genomics Facility (GEFA) with special thanks to Yeraldin Chiquinquira Castillo De Spelorzi and Edoardo Henzen. A special thanks to Andrea Contestabile for Neurod1 antibody. This work was partially funded by Project “National Center for Gene Therapy and Drug based on RNA Technology” (CN00000041). Financed by NextGenerationEU PNRR MUR – M4C2 – Action 1.4 - Call “Potenziamento strutture di ricerca e di campioni nazionali di R&S” (CUP J33C22001130001) to SG and by the Marie Curie MINDED grant 754490 to AS. We acknowledge that this research was conducted using the IIT HPC infrastructure.

## AUTHOR CONTRIBUTIONS

Conceptualization: AS, DD, RS, SG; methodology: AS, FA, DD, LS, DV, LP; investigation: AS, FA, DD, LS, DM, PL, DV, LP; visualization: AS; funding acquisition: SG, RS; project administration: SG, RS; supervision: SG, RS; writing – original draft: AS, FA, SG, RS; writing – review & editing: All authors.

---

**Fig. S1.**
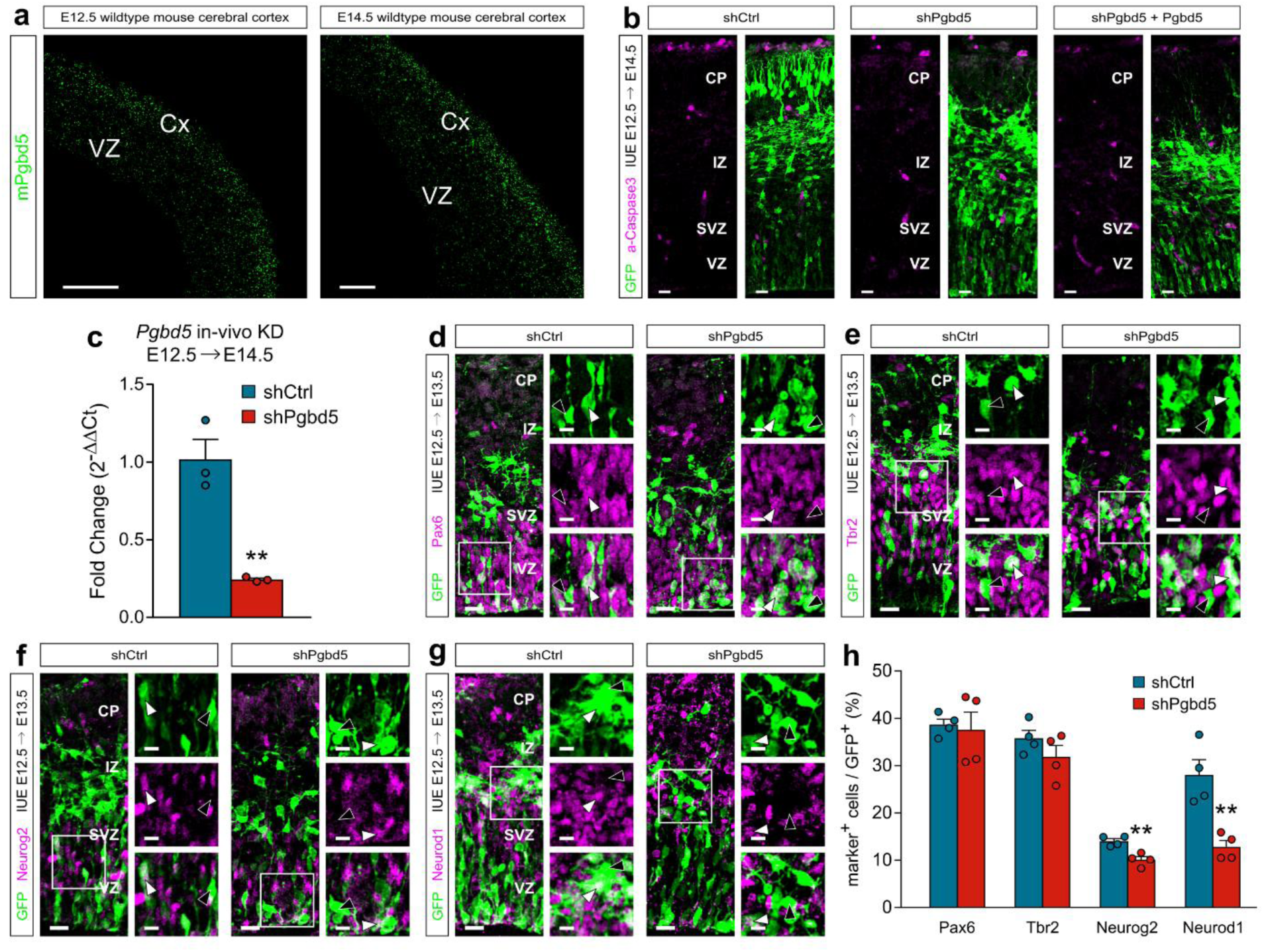
Pgbd5 is expressed in embryonic mouse neocortex and is involved in neurons generation. **a**, *Pgbd5 in situ* hybridization on E12.5 (left) and E14.5 (right) mouse coronal cerebral cortices. Scale bar 100um. **b**, IF staining of cleaved Caspase 3 apoptosis marker. Scale bar 20um. **c**, Expression of *Pgbd5* RNA in GFP^+^ cells from E14.5 cortices injected with shCtrl or shPgbd5 at E12.5. RNA levels are normalized on shCtrl group. n = 3 technical replicates. Unpaired Student’s t-test. **d-g**, IF staining for NPCs cells (Pax6, Tbr2) and pro-neuronal (Neurog2, Neurod1) markers at E13.5, shRNAs *iue* at E12.5. Scale bar 20um, close-up scale bar 10um. **h**, Quantification of marker^+^/GFP^+^ cells at E13.5 in E12.5 shPgbd5- vs shCtrl-injected brains. Data are mean ± s.e.m.; *n* = 4 each. Unpaired Student’s t-test. Cx, cerebral cortex; VZ, ventricular zone; SVZ, subventricular zone; IZ, Intermediate zone; CP, cortical plate. \*\**P* < 0.01.

**Fig. S2.**
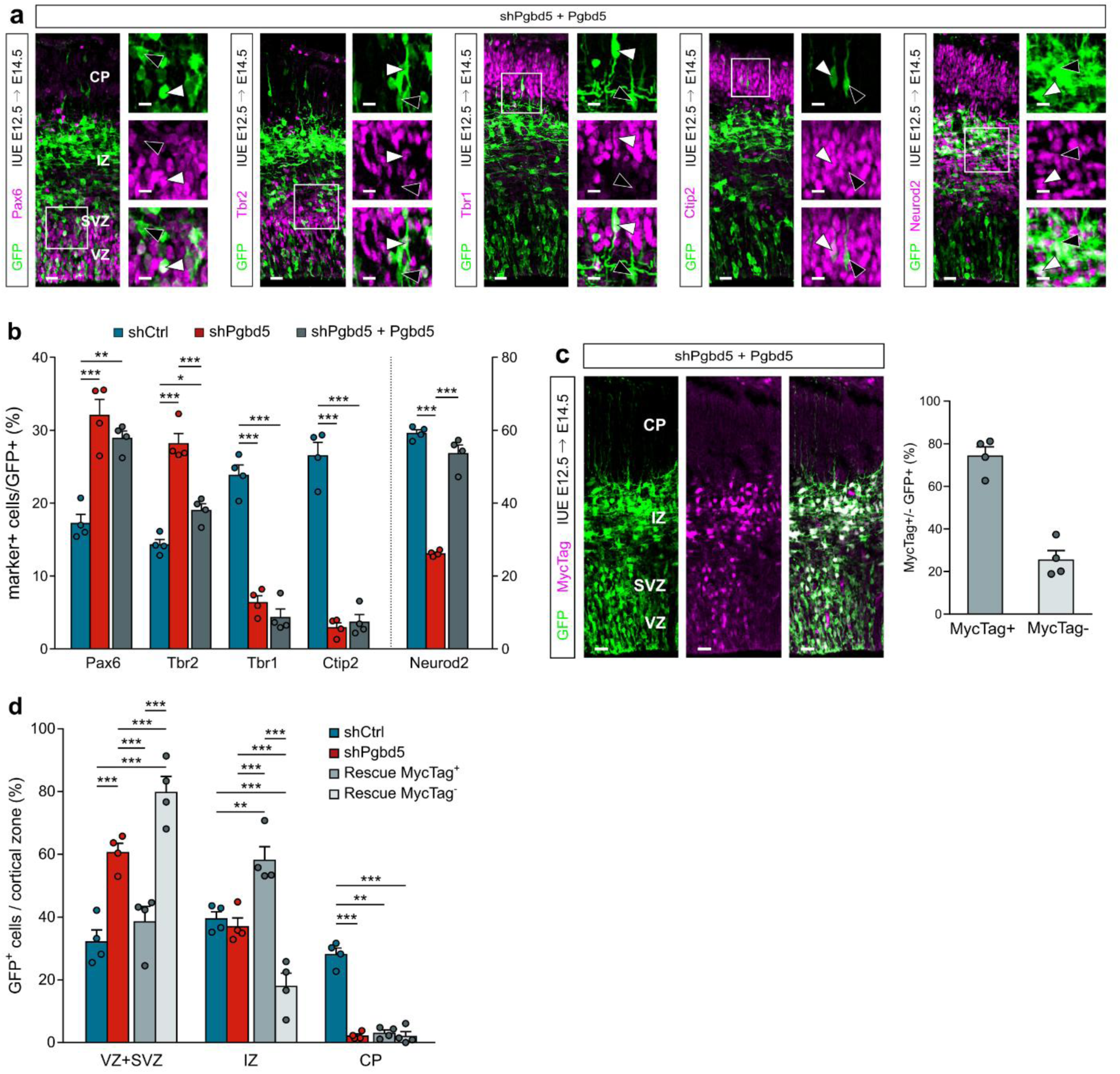
Markers expression and distribution of Pgbd5-rescued cells. **a**, IF staining for NPCs (Pax6, Tbr2), neuronal cells (Neurod2) and DL neurons (Tbr1, Ctip2) markers at E14.5 in *Pgbd5*-rescued cells at E14.5. Positive expression (white arrowheads) vs. no expression (black arrowheads). Scale bar 20um, close-up scale bar 10um. **b**, Quantification of marker^+^/GFP^+^ cells at E14.5 in rescue experiments. Data are mean ± s.e.m.; n = 4 each. Two-way ANOVA with Tukey’s multiple comparisons test. **c**, E14.5 IF staining and quantification of MycTag^+/-^ GFP^+^ cells in *Pgbd5* rescue experiments. Scale bar 20um. **D**, E14.5 MycTag^+/-^ GFP^+^ cells distribution in VZ+SVZ, IZ and CP of E12.5 shCtrl- vs. shPgbd5- vs. *Pgbd5*-rescued cells. Data are mean ± s.e.m.; n = 4 each. Two-way ANOVA with Tukey’s multiple comparisons test. VZ, ventricular zone; SVZ, subventricular zone; IZ, Intermediate zone; CP, cortical plate. \**P* < 0.05, \*\**P* < 0.01, \*\*\**P* < 0.001.

**Fig S3.**
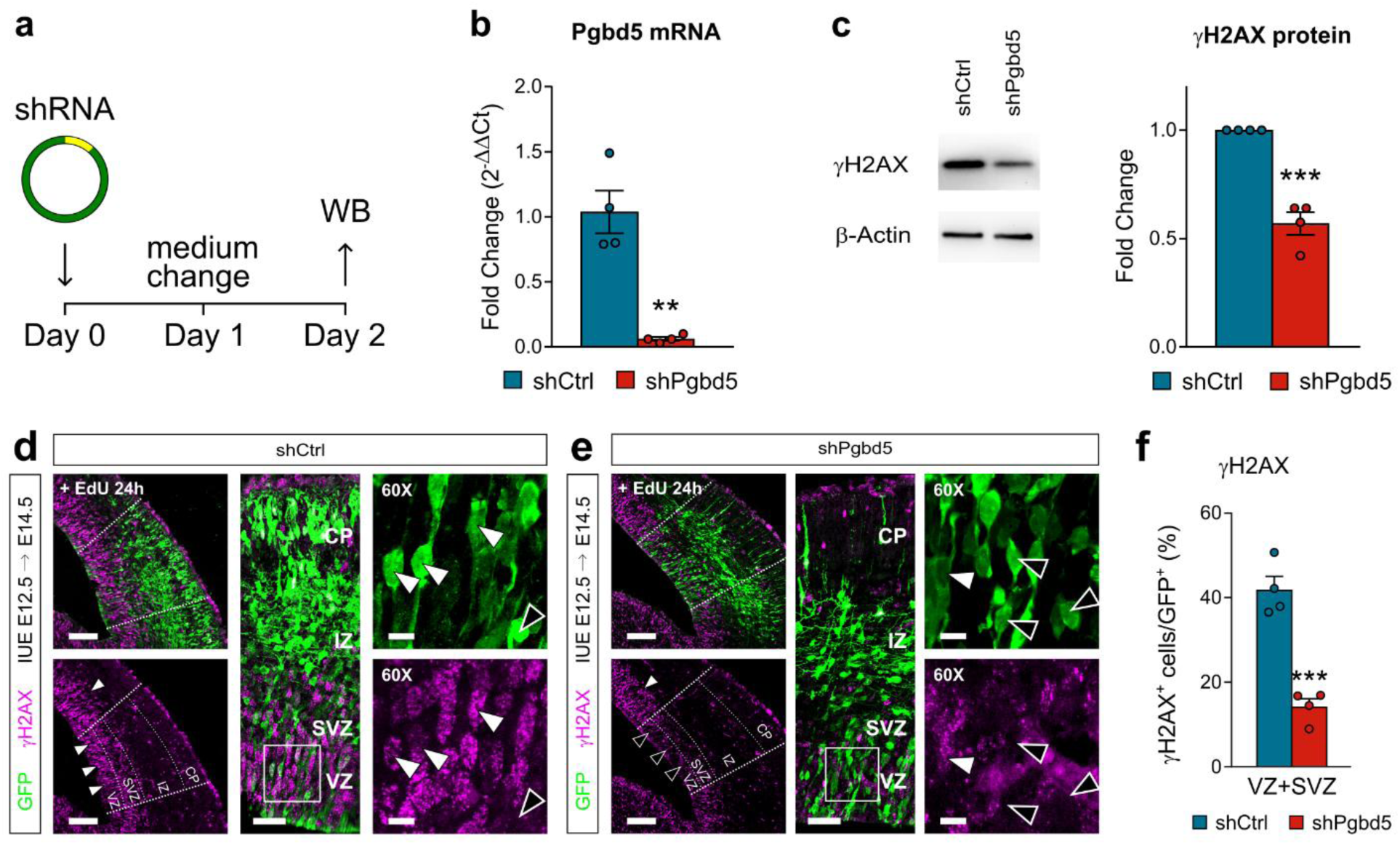
Pgbd5 knockdown affects physiological and induced occurrence of DNA double-strand breaks in cells. **a**, Schematic of in-vitro *Pgbd5* manipulation in N2a cell cultures. **b**, Expression levels of *Pgbd5* in shCtrl- and shPgbd5-treated N2a cell cultures. RNA levels are normalized on shCtrl group. n = 4 independent experiments. Unpaired Student’s t-test. **c**, Protein expression analysis. One representative experiment is shown (left). Band intensity was normalized to the relative β-actin band. Fold change values were calculated normalizing on shCtrl cells. Data are mean ± s.e.m.; *n* = 4 independent experiments. Unpaired Student’s t-test. **d-e**, γH2AX IF signal in shCtrl- (d) vs. shPgbd5-injected brains (e). Positive expression (white arrowheads) vs. no expression (black arrowheads). Scale bar iue area 100um, scale bar 40um, close-up scale bar 10um. **f**, Quantification of γH2AX^+^/GFP^+^ cells at E14.5 + EdU 24h. Data are mean ± s.e.m.; *n* = 4 shCtrl, *n* = 4 shPgbd5. Unpaired Student’s t-test. VZ, ventricular zone; SVZ, subventricular zone; IZ, Intermediate zone; CP, cortical plate. \*\**P* < 0.01, \*\*\**P* < 0.001.

**Table S1.** (separate file). See file Table_S1_DEgenes.xlsx

**Table S2.** (Separate file) See file Table_S2_GO.xlsx **Table S3**

**Table S3.** Antibodies table.

| Ab name | Host species | Company | Catalog number | Dilution | Application |
| --- | --- | --- | --- | --- | --- |
| anti-GFP | chicken polyclonal | Abcam | ab13970 | 1:1000 | IF |
| anti-Ctip2 | rat monoclonal | Abcam | ab18465 | 1:1000 | IF |
| anti-Tbr1 | rabbit polyclonal | Abcam | ab31940 | 1:1000 | IF |
| anti-Tbr2 | rabbit monoclonal | Abcam | ab183991 | 1:1000 | IF |
| anti-Pax6 | rabbit polyclonal | Millipore | AB2237 | 1:1000 | IF |
| anti-NeuroD1 | goat polyclonal | Santa Cruz Biotech. | sc-1084 | 1:200 | IF |
| anti-Neurog2 | rabbit polyclonal | Cell Signaling | 13144 | 1:500 | IF |
| anti-Cleaved Caspase-3 | rabbit monoclonal | Cell Signaling | 9664 | 1:500 | IF |
| anti-gH2AX | rabbit polyclonal | Cell Signaling | 9718S | 1:500 | IF / WB |
| AlxFluor-488 anti-Chk | goat | Thermo Scientific | A11040 | 1:1000 | IF |
| AlxFluor-546 anti-Rb | goat | Thermo Scientific | A-11035 | 1:1000 | IF |
| AlxFluor-647 anti-Rb | goat | Thermo Scientific | A-21244 | 1:1000 | IF |
| AlxFluor-647 anti-Rat | goat | Thermo Scientific | A-21247 | 1:1000 | IF |
| AlxFluor-546 anti-Goat | donkey | Thermo Scientific | A-11056 | 1:1000 | IF |
| AlxFluor-647 anti-Rb | donkey | Thermo Scientific | A-21447 | 1:1000 | IF |
| anti-b-Actin | mouse monoclonal | Merck | A3854 | 1:10000 | WB |

**Table S4.**
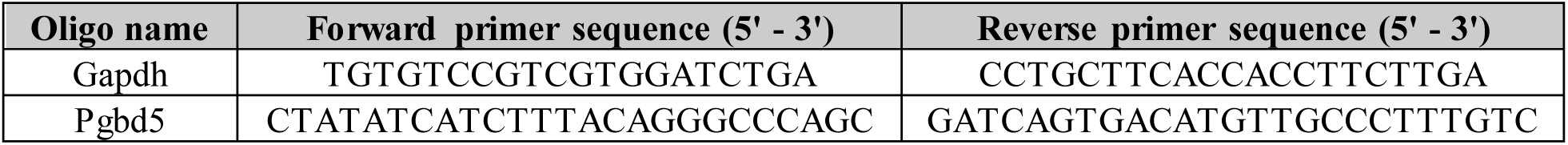
RT-qPCR primer table.

**Table S5.**
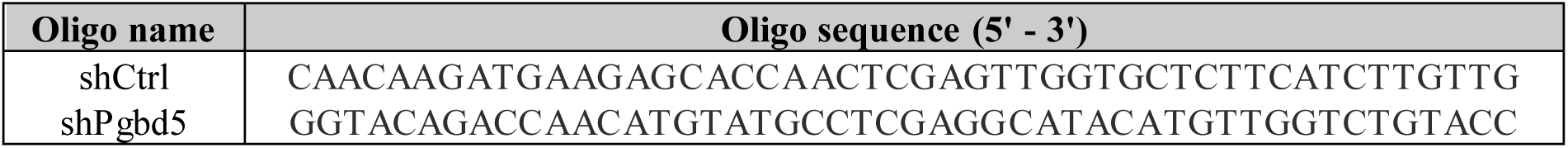
shRNA sequences.

